# A diallel of the mouse Collaborative Cross founders reveals strong strain-specific maternal effects on litter size

**DOI:** 10.1101/458877

**Authors:** John R. Shorter, Paul L. Maurizio, Timothy A. Bell, Ginger D. Shaw, Darla R. Miller, Terry J. Gooch, Jason S. Spence, Leonard McMillan, William Valdar, Fernando Pardo-Manuel de Villena

## Abstract

Reproductive success in the eight founder strains of the Collaborative Cross (CC) was measured using a diallel-mating scheme. Over a 48-month period we generated 4,448 litters, and 24,782 weaned pups were used across 16 different published experiments. We identified factors that affect the average litter size in a cross by estimating the overall contribution of parent-of-origin, heterosis, inbred, and epistatic effects using a Bayesian zero-truncated overdispersed Poisson mixed model. The phenotypic variance of litter size has a substantial contribution (79%) from unexplained and environmental sources, but no detectable effect of seasonality. Most of the explained variance was due to additive effects (9.2%) and parental sex (maternal vs paternal strain; 5.8%), with epistasis accounting for 3.4%. Within the parental effects, the effect of the dam’s strain explained more than the sire’s strain (13.2% vs. 1.8%), and the dam’s strain effects account for 74.2% of total variation explained. Dams from strains C57BL/6J and NOD/ShiLtJ increased the expected litter size by a mean of 1.66 and 1.79 pups, whereas dams from strains WSB/EiJ, PWK/PhJ, and CAST/EiJ reduced expected litter size by a mean of 1.51, 0.81, and 0.90 pups. Finally, there was no strong evidence for strain-specific effects on sex ratio distortion. Overall, these results demonstrate that strains vary substantially in their reproductive ability depending on their genetic background and that litter size is largely determined by dam.strain rather than sire.strain effects, as expected. This analysis adds to our understanding of factors that influence litter size in mammals, and also helps to explain breeding successes and failures in the extinct lines and surviving CC strains.

## Introduction

A critical part of fitness is optimizing the balance between numbers of offspring, offspring body size, and reproductive timing, leading to strong selection for these traits (Smith and Fretwell 1974; Beauchamp and Kacelnik 1990). Fundamental fitness traits, like litter size (*i.e.,* the number of pups born per litter), should have low contributions of genetic variance due to rapid fixation of alleles that increase fitness (Mousseau and Roff 1987; Price and Schluter 1991; Merilä and Sheldon 1999; Stirling *et al.* 2002; Peripato *et al.* 2004). Estimates of litter size heritability vary from 5% to 20% in mouse, rabbit, and porcine populations (Falconer 1960; Avalos and Smith 1987; Falconer and Mackay 1996; Johnson *et al.* 1999; Garcia and Baselga 2002; Peripato *et al.* 2004; Gutiérrez *et al.* 2006) leading to the majority of variation in litter size being non-heritable. Non-heritable factors include seasonality, food availability, paternal and maternal effects, dominance and epistasis, as well as unknown factors. Additionally, sexual conflict over parental investment may exist between females and males, and this in turn could affect litter size (Trivers 1974; Hager and Johnstone 2003). These factors make it difficult to obtain clear measurements of litter size in wild populations. A controlled laboratory environment is therefore necessary to measure genetic effects of litter size (Jinks and Broadhurst 1963; Roberts 1960; Bandy and Eisen 1984; Hoornbeek 1968; Peripato *et al.* 2004; Gutiérrez *et al.* 2006; Varona and Sorensen 2014).

The Collaborative Cross (CC) and its eight founder strains are an important resource for studying complex traits, establishing mouse models for human disease, and understanding the mouse Diversity Outbred (DO), which originated from the CC (Ferris *et al.* 2013; Chesler 2014; Rogala *et al.* 2014; Rasmussen *et al.* 2014; Gralinski *et al.* 2015, 2017; Schoenrock *et al.* 2017; Maurizio *et al.* 2018). The CC resource and its founder strains can also be used to study reproductive ability, which is especially interesting because this population has an established record both of breeding successes and of failures. For example, litter size in the early CC lines shows a steady decline over the first six generations of inbreeding (Philip *et al.* 2011). In addition, nearly 95% of all CC lines have become extinct, primarily due to subspecific genomic incompatibilities (Shorter *et al.* 2017). Although standard reproductive phenotypes have been measured, mostly in male CC founders (Odet *et al.* 2015), the genetic and non-genetic control over CC breeding success has never been thoroughly characterized, and is likely to contain vital information that may be used to improve CC breeding in the future.

Given the growing popularity of the CC as a community resource, we investigated reproductive ability across the eight founders of the CC to better understand the genetic and nongenetic factors that affect their fertility and breeding. Using an 8 × 8 diallel design, we measured weaned litter sizes from over 4400 litters arising from 62 crosses across four years of breeding. Adapting a recently developed statistical model of diallel effects (Lenarcic *et al.* 2012), we quantified both the genetic contributions that shape litter size and the contributions of several environmental factors. Our results provide a detailed account of breeding patterns across and between the eight founders. This experiment also informs us about genetic combinations that are highly or marginally productive in the CC, helping us to better understand CC fertility problems and line extinction.

## Materials and Methods

The mouse inbred strains used in these experiments are the eight founder strains of the Collaborative Cross (CC) (Collaborative Cross Consortium 2012). The founders of the CC include five classical strains, A/J (AJ), C57BL/6J (B6), 129S1/SvImJ (129S1), NOD/ShiLtJ (NOD), NZO/HlLtJ (NZO), and three wild-derived strains, CAST/EiJ (CAST), PWK/PhJ (PWK), and WSB/EiJ (WSB). Mice originated from a colony maintained by Gary Churchill at Jackson Laboratory, and were transferred to the FMPV lab at the University of North Carolina (UNC) in 2008. The original colony also produced most of the G1 breeders that populated the inbred funnels at ORNL, TAU and Geniad (Srivastava *et al.* 2017). All mice described here were reared by the FPMV lab at UNC. Mice were bred at the UNC Hillsborough vivarium from 2008-2010 and bred at the UNC Genetic Medicine Building (GMB) vivarium from 2010-2012. A total of 4,448 litters resulting from crosses between 1,478 individual dams and 1,238 individual sires were born from all eight inbred crosses and 54 of 56 reciprocal F1 hybrid crosses, excluding hybrids between NZO × CAST and NZO × PWK, which are known unproductive crosses (Chesler et al. 2008). The directions of all crosses are described as female by male (i.e., dam.strain × sire.strain), unless otherwise noted. Litter size and sex were determined at weaning by visual inspection. Animals were kept on a 14-hour, 10-hour light/dark schedule with lights turned on at 7:00 AM; temperature was maintained at 20°-24° with relative humidity between 40-50%. Mice were housed in standard 20×30-cm ventilated polysulfone cages with standard laboratory grade Bed-O-Cob bedding. Water and Purina Prolab RMH3000 were available *ad libitum.* Mouse chow was supplemented with Fenbendazole (Feb 2010) two weeks before and two weeks after transportation to the GMB facility to eliminate possible pinworms. Selemectin treatment was dropped onto the coats of mice before transfer to remove mites from the cages. These treated cages were not opened until after their arrival at UNC GMB. All animal rearing and breeding was conducted in strict compliance with the *Guide for the Care and Use of Laboratory Animals* (Institute of Laboratory Animals Resources, National Research Council 1996, https://www.ncbi.nlm.nih.gov/books/NBK232589/). The Institutional Animal Care and Use Committee of the University of North Carolina approved all animal use and research described here.

### Statistical Analysis

#### Testing environmental interactions

Significant environmental interactions were determined using ANOVA, using JMP 12 software (JMP®, Version 12. SAS Institute Inc., Cary, NC, 1989-2007). Tested effects included a season effect (average weaned litter size in each month over every year), non-seasonal factors using a year-by-month effect (average weaned litter size for all months across all years), and a litter order effect (average weaned litter size in each subsequent dam litter). We performed the ANOVA on litter size counts from the eight inbred matings only, because of their robust sample sizes throughout the four years of breeding.

#### Diallel analysis of litter size

To estimate the overall contributions of heritable factors affecting litter size in our population, we adapted a previously published linear mixed model, BayesDiallel (Lenarcic et al. 2012), which performs this estimation for continuous phenotypes, to the setting of a discrete count-based phenotype. To do this we reimplemented the BayesDiallel Gibbs sampler as a generalized linear mixed model (GLMM) using the R software package MCMCglmm (Hadfield 2010). Let *y_i_* be the number of pups born to litter *i*, where 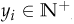, and denote the corresponding pair of parental strains, namely maternal strain *j* and paternal strain *k*, using the notation {*jk*} [*i*]. The effect of parental strains on y_i_ was modeled using an overdispersed zero-truncated Poisson (ZTP) regression (data scales of the model in brackets):

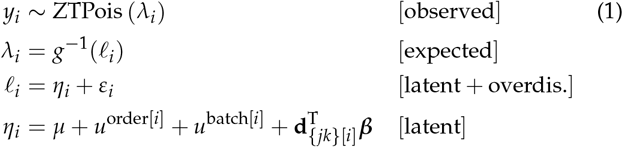

where ZTPois(*λ_i_*) denotes a Poisson distribution with expected value *λ_i_* but conditional on having observed that *y_i_* ≠ 0 (**Appendix A**), g is the link function *g(x)* = log(x), with inverse *g ^−1^(x) = e^x^*, that relates *λ_i_* to a latent scale *ℓ_i_*, and *η_i_* is a linear predictor on that latent scale with an error term 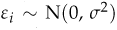 providing overdispersion. The linear predictor *η_i_* is then composed of an intercept μ, random effects for litter order (*i.e.,* parity) *u*^order[i]^ and batch (month) *u*^batch[i]^ respectively (detailed below), where for example batch *[i]* refers to the batch containing litter *i*, and a further linear predictor d^T^*_{jk}[i]_ β* (detailed below) that specifies the contributions of parental strains *j* and *k* (detailed below). The random effects for litter order and batch were both modeled as categorical random intercepts, *u*^batch^ ∼ N(0, *τ*^2^_batch_) and *u*^order^ ∼ N(*c*^order^ · *α, τ*^2^_order_), where litter order is additionally predicted by *c*^order^, the order as a number, with fixed effect *α.*

The parental strain contribution d^T^_{jk}_*β*, is equivalent to the ‘fullu’ (full, unsexed) model described in (Lenarcic *et al.* 2012), and is given below:

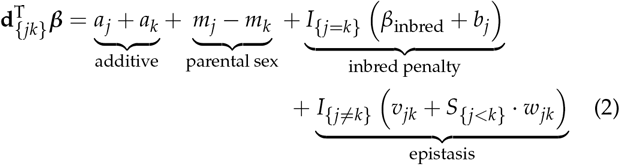

where I_{A}_ is an indicator and equal to 1 if A is true and 0 otherwise, S_{A}_ is a sign variable equal to 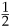 if A is true and 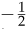 otherwise, β_inbred_ is a fixed effect of being inbred, and all other variables (in classes *a, m, b, v, w*) are modeled as random effects with class-specific variances, for example, a1,…,a8 are modeled as a_j_ ∼ N(0, τ^2^_a_). The *a* (“additive”) class represents strain-specific dosage effects, the *m* (“parental sex”) class represents asymmetric parent-of-origin effects, where positive values indicate that maternal effects are greater than paternal effects for a given strain, inbred penalty effects are comprised of overall (fixed) and strain-specific (random) b parameters, and epistatic effects are strain-pair specific deviations, both symmetric (*v*) and asymmetric (*w*).

To provide directional parent-of-origin effects for maternal and paternal strain we defined, by reparameterization of the additive and parental sex effect, the following two additional types of effect:

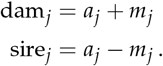

For example, dam_B6_ = *a*_B6_ + *m*_B6_ is the B6-specific dam (maternal strain) effect and sire_B6_ = *a*_B6_ - *m*_B6_ is the B6-specific sire (paternal strain) effect. Posterior samples of these quantities were obtained as a post-processing step by reparameterizing posterior samples of *a_j_* and *m_j_*. Obtaining these as a post-processing step, rather than explicit modification of Equation 2, preserves the original BayesDiallel model and therefore has no effect on the effect estimates (or variance projection estimates) for the other diallel categories. (Parameter definitions summarized in Table S1.)

Priors were chosen to be minimally informative. For fixed effects (*μ, α*, and *β*_inbred_) we used N (0,1 x 10^3^). For variances of random effects 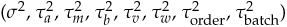 we used an in verse gamma distribution with scale and shape both equal to 0.001. Posterior effect estimates are presented as posterior mean (and median), and the 95% highest posterior density (HPD) interval.

In order to stably estimate the contribution of each variance class to the overall phenotype, we used the diallel variance projection [VarP; Crowley et al. (2014)]. This is a heritability-like measure that partitions the overall phenotypic variance of an idealized future diallel experiment into additive, parent-of-origin (parental sex), inbred (dominance), epistatic, and other random/fixed effects categories in the diallel. VarPs were calculated both from the ZTP model described above, and, for comparison, from the standard Gaussian-outcome BayesDiallel model (using the BayesDiallel software) (**Appendix A**). The standard BayesDiallel model was applied to our data after litter size was subject to a variance-stabilizing transformation, the square root, this corresponding to the linear mixed model approximation to the ZTP,

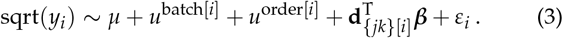

The standard BayesDiallel model under this transformation was also used to evaluate evidence in our data set for the inclusion (or exclusion) of each variable or class of variables in improving the model fit. Specifically, we calculated for each variance class a posterior model inclusion probability (MIP) Lenarcic *et al.* (2012). MIPs with a value of 1 indicate strong evidence to retain a variable or inheritance class in the model, whereas MIPs near 0 indicate strong evidence for exclusion of a variable/class from the model. MIPs within the range of (0.05,0.25] or [0.75,0.95) indicate positive evidence, (0.01,0.05] or [0.95,0.99) indicate strong evidence, and [0,0.01] or [0.99,1] indicate very strong evidence for model inclusion, approximately following the scheme of Kass and Raftery (1995).

#### Diallel analysis of sex ratio

To model diallel effects on sex ratio we recast the BayesDiallel model as a binomial GLMM. Letting *y_i_* be the number of males out of a total of *n_i_* pups for litter *i*, we model where *π_i_* is the expected proportion of males predicted for litter *i*, the link function is g(x) = logit(x) = log(x) / (1 — x), with inverse link *g^−1^* (x) = logit^−1^ (x) = expit(x) = *e*^x^/ (1 + *e*^x^), and *ℓi*, which represents *π_i_* on the latent scale, is modeled using the BayesDiallel hierarchy as in Equation 1

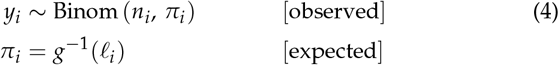

Additional details about our statistical modeling approaches are provided in **Appendix A** (litter size) and **Appendix B** (sex ratio).

### Data Availability

File S1 contains all breeding data used for the analysis in this study. File S2 contains the scripts and software used for the analysis in this study. File S3 contains the Supplemental Figures and Tables. The data and code used to analyze the data are available at https://github.com/mauriziopaul/litterDiallel, with a static version available at https://doi.org/10.5281/zenodo.1435829. Phenotype data from this study is available on the Mouse Phenome Database (Bogue *et al.* 2018) at https://phenome.jax.org/projects/Shorter with persistent identifier RRID:SCR003212, Accession number MPD:623, project Shorter3.

## Results

### Litter size is affected by housing facility but not season

We bred litters in an 8x 8 inbred diallel of the CC founder strains, and generated all eight inbreds and 54 of 56 possible reciprocal F1 hybrids (Figure 1, Figure S1-S3). The eight inbred strains, displayed across the diagonal, were mated at higher frequencies both for maintenance of inbred strains and propagation of the diallel. For all 62 genetic inbred and hybrid crosses, we recorded the following information: mated pairs, wean dates, litter size at weaning, including total and sex-specific counts (File S1). This diallel cross was originally designed and maintained for the generation of F1 mice for several experimental projects (Koturbash *et al.* 2011; Aylor *et al.* 2011; Mathes *et al.* 2011; Kelada *et al.* 2012; Didion *et al.* 2012; Collaborative Cross Consortium 2012; Calaway *et al.* 2013; Crowley *et al.* 2014; Phillippi *et al.* 2014; Odet *et al.* 2015; Crowley *et al.* 2015; Morgan *et al.* 2016; Percival *et al.* 2016; Shorter *et al.* 2017; Oreper *et al.* 2017; Maurizio *et al.* 2018). As a result, certain reproductive measurements such as time between litters and maximum number of offspring per cross were necessarily biased by experimental breeding requirements. Average weaned litter size, however, is a reproductive trait that should be well-estimated independently of these factors.

**Figure 1.**
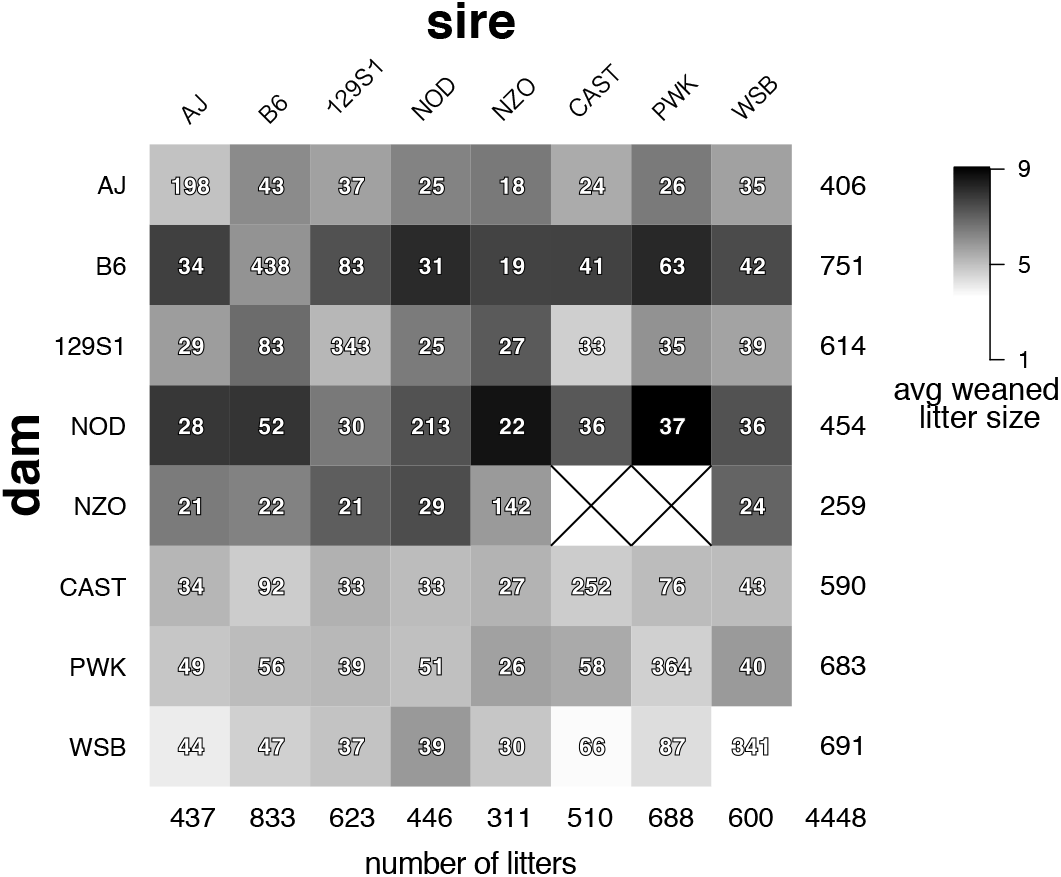
Diallel crossing scheme and weaned pup distribution. The number of litters observed per cross is given by the integers, with the largest sample sizes, along the diagonal, corresponding to the production of inbred parental strains. Column and row sums are given along the bottom row and rightmost column, respectively. A total of 4,448 litters were evaluated for this analysis, resulting in a total of 24,782 weaned pups. The shading within each box corresponds to the average number of weaned pups per litter in each cross, with averages ranging from 3.6 to 9.1 pups per litter. Litters for which no pups survived until weaning were not included in our analysis. The symbol “×” is used to indicate incompatible crosses that do not produce any litters.

We measured litter size and report the mean number of weaned pups per litter for the 62 viable crosses in the diallel (Figure 1). A wide distribution of litter sizes was observed, ranging from an average of 3.6 weaned pups for WSB × WSB crosses, to an average of 9.1 weaned pups for NOD × PWK crosses, with an overall mean of 5.5 weaned pups per litter. Examining average litter sizes of the inbred strains across all 4 years, we found neglible evidence of consistent seasonal effects (f_11,2283_ = 1.272 *p* = 0.23) (Figure S4) but positive evidence of non-seasonal patterns: a modeled ‘year.month’ covariate significantly affected litter size (f_47,2247_ = 2.44 *p* < .0001; see also Figure S5). The non-seasonal effect could be driven by the relocation of mice between vivariums, which occurred in February 2010. The housing facility may have an impact on litter size as well. To test this, we measured average litter size differences between the two housing facilities and found that two founder strains, 129S1 and WSB, had significantly larger average litter sizes at the Hillsborough facility than at UNC GMB with a difference of 5.87 to 4.42 weaned pups per litter (p < 0.0001) for 129S1, and 3.84 to 3.42 weaned pups per litter (p = 0.029) for WSB. The six other founder strains did not significantly differ in their average litter sizes between the two facilities, but tended to have smaller weaned litters at UNC GMB (see Discussion). For average litter size across the diallel, we tested for an effect of litter number (birth parity number, for a given mating pair) on the number of pups weaned in each litter. Previous research suggests that the first litter can be significantly smaller than subsequent litters due to various biological factors (De la Fuente and San Primitivo 1985). This effect was consistent in the diallel: we observed that there was significant reduction in litter size in the first litter (*p* < 0.0001) compared to the overall linear effect that parity has on reducing litter size (Figure S5).

### Litter size is moderately heritable, and maternal effects account for the majority of explained variation

To estimate founder strain effects on litter size, we used a Bayesian regression model that decomposes the phenotypic variation in a diallel design into genetic and parent-of-origin contributions (Lenarcic *et al.* 2012). Using this model, litter size was estimated to have narrow sense heritability (*h*^2^ approximated by VarP[additive]) accounting for 9.18% of the variance, with parent-of-origin effects (VarP[parental.sex]) accounting for 5.77%, the fact of being inbred (VarP[inbred.overall]) at 1.43%, and strain-by-strain interactions (VarP[epistatic.symmetric] + VarP[asymmetric.epistatic]) at 3.40% (Figure 2A); in aggregate, these factors explained 17.73% of the total variance for litter size in the founder diallel.

**Figure 2.**
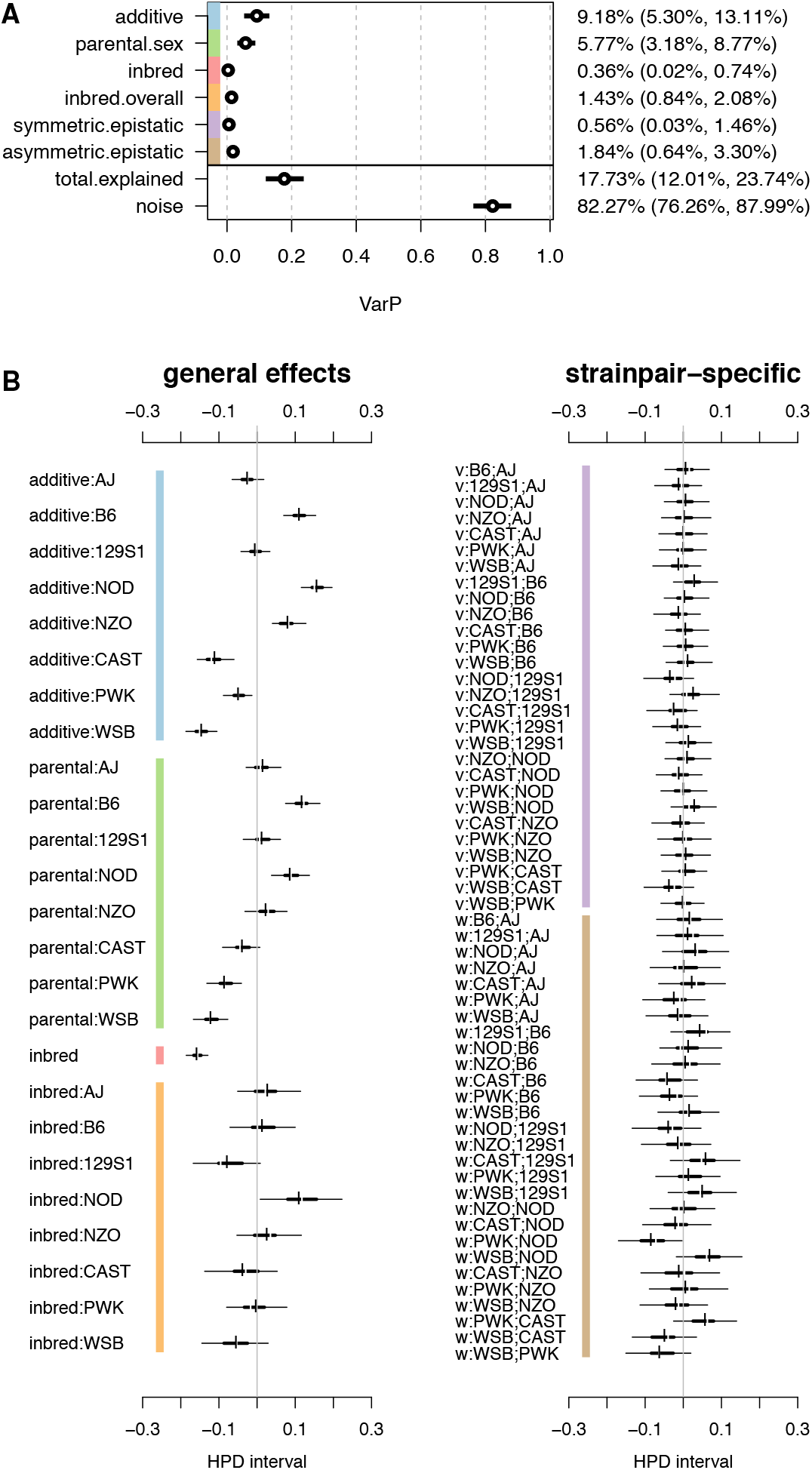
Diallel effects and variance contributions for weaned litter size. (A) Variance contributions of distinct effect classes, reported as posterior means and 95% HPDs of variance projections (VarPs) on weaned litter size. (B) Diallel effects, including (left) strain-specific additive, parental sex, and inbred effects, and (right) epistatic effects between each pairwise cross. For each parameter, thin and thick horizontal lines represent 95% and 50% highest posterior density (HPD) intervals of effects, respectively, and vertical break and dash give posterior median and mean, respectively. The effects are in relation to an overall mean litter size of 5.46 (95% HPD: 5.00-6.10). The gray vertical lines indicate zero. Effects are shown as the log, or latent, scale effects on the mean litter size attributable to each strain or strainpair and inheritance group, where values are centered at 0 for each random effect class. Intervals that exclude zero have non-negligible effects on the mean litter size. Labels with “v” or “w” refer to symmetric or asymmetric epistatic effects, respectively. Colored bars indicate corresponding variance classes in (A) and (B).

In more detail, we present estimates for all modeled diallel effects as posterior means and highest posterior density (HPD) intervals (Figure 2B). Parameters are divided into two groups: general effects and strain pair-specific effects. General effects comprise strain-specific additive effects (additive), strain-specific and overall inbred (inbred), and strain-specific parent-of-origin (parental sex) effects. Strain pair-specific (epistatic) effects are the effects that arise specifically in crosses of two heterologous strains, with ‘v’ referring to symmetric epistatic and ‘w’ referring to asymmetric epistatic effects (pairwise parent-of-origin effects). All classes of effect were determined to add positive value to the model fit (either strong or very strong evidence for inclusion; MIP > 0.986) except parental sex (the difference between male and female), whose value was equivocal (MIP=0.34) Table S2).

Under the general effects, we see significant positive additive effects on average litter size from B6, NOD, and NZO and significant negative additive effects on litter size from CAST, PWK, and WSB strain dosages. A similar pattern is seen in the parental sex effects, where B6 and NOD dosages have a significant positive effect on average litter size, whereas CAST, PWK, and WSB have significant negative effects. The “inbred” effect is negative indicating that inbred status has a negative overall effect on average litter size, regardless of parental strain. Each strain also has an individual inbred effect in addition to an overall inbred effect; the only strain with a positive inbred effect is NOD. When the individual inbred effects are taken into account with the overall “inbred” effect, inbred litters are on average slightly smaller than their heterozygous counterparts. For strain pair-specific epistatic effects, there are a few marginally noteworthy effects, with the most prominent being a negative asymmetric epistatic effect for PWK × NOD.

To more clearly differentiate the contributions of the mother vs. the father strain, we reparameterized the BayesDiallel model to capture effects specific to dam.strain and sire.strain (Figure 3). B6 and NOD dams increase litter sizes by more than 1.26 fold, by an average of 1.66 and 1.79 pups, respectively, regardless of sire. CAST, PWK, and WSB dams tend to decrease average litter size by 0.90, 0.81, and 1.51 pups. The sire effect is similar, with NOD and NZO sires having larger litters and CAST sires producing smaller litters. As expected, we see that the dam.strain has a much larger influence on the variation of litter size compared with the sire.strain (13.15% vs. 1.81%).

**Figure 3.**
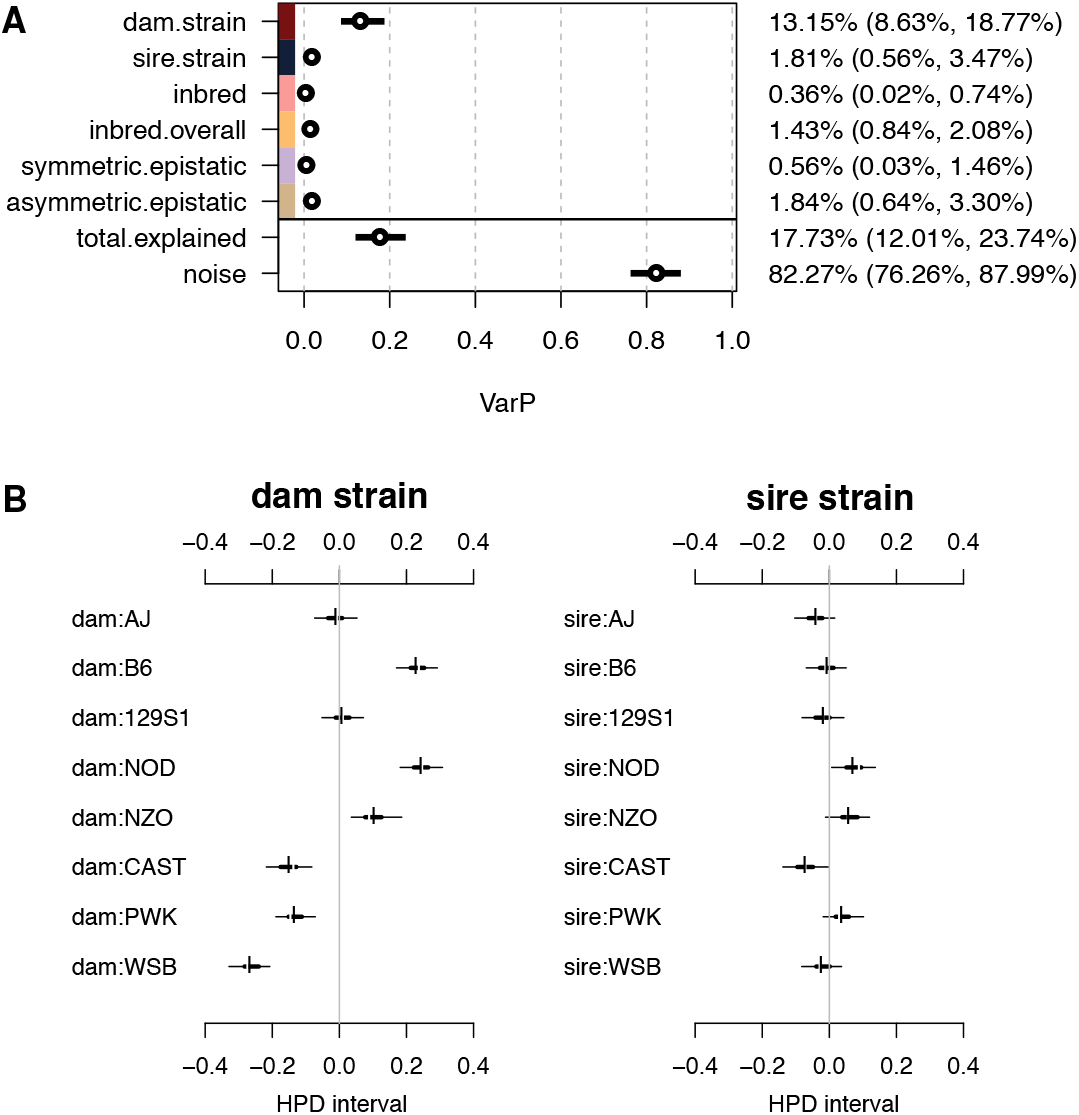
Dam.strain and sire.strain variance contributions and estimates of effects on weaned litter size. These effects are a reparameterization of additive and parental.sex effects from the previous analysis. (A) Variance contributions of distinct effect classes, reported as posterior means and 95% HPDs of variance projections (VarPs) on weaned litter size. (B) Estimates for the maternal (“dam.strain”) and paternal (“sire.strain”) effects on litter size, as calculated from the additive and parental sex parameters in Figure 2, with HPD intervals defined correspondingly.

### No significant strain effects on sex ratio

We examined genetic and non-genetic effects on the average sex ratio per litter and found no evidence that sex ratio was skewed (Figure S6;Figure S7). The overall mean for sex bias, quantified as number of male pups weaned divided by the total number of weaned pups, was 0.4979, and did not significantly differ from our expectation of a 50:50 sex ratio (Binominal test, two-tailed *p* = 0.219). For the eight inbred founders, 129S1 is the only strain that departed significantly from expectation, with slight reduction of males 0.475:0.525 (Binominal test, two-tailed *p* = 0.041). However, correction for multiple testing shows no significant sex ratio bias of any inbred strain. The outbred crosses have substantially fewer litters and offspring than the inbred matings, leading to less balanced sex ratios; however, when multiple testing is accounted for, they also show no significant deviations from expected sex ratios.

## Discussion

We have investigated factors influencing litter size in the eight CC founders and their F1 hybrids. We note that litter size is a component of reproductive performance, but distinct from total strain productivity; a study on total strain productivity would need to take into account litter size, numbers of litters, maximum reproductive age, and pup survival until breeding. Our results illustrate how mammalian litter size in a full diallel design is influenced by genotypic and environmental variation. They also address some of the factors that contributed to line extinction and breeding problems in generating the CC (Chesler *et al.* 2008; Philip *et al.* 2011; Collaborative Cross Consortium 2012; Shorter *et al.* 2017).

We estimated that genetic (additive, inbred, and epistatic) effects on average litter size explained 17.73% of variation, suggesting that most of the phenotypic variation arises from unexplained environmental effects. Within the explained variance, our estimate of the narrow sense heritability (*h*^2^ = 0.092) for litter size may seem small, but it is within the range uncovered in previous research (Falconer 1960; Peripato *et al.* 2004; Gutiérrez *et al.* 2006). Compared with the overall average litter, we observed substantial positive effects of B6 and NOD strains, from both additive genetic and parent-of-origin parameters, and substantial negative effects of PWK and WSB (Figure 2). We also observed, as expected, that lower litter size was associated with being inbred (Figure 2A).

During the G1 and G2 out-crossing generations of the CC, mean litter size was lower for crosses involving wild-derived strains, CAST, PWK and WSB (Philip *et al.* 2011). A similar pattern is observed here, but these effects are determined to be specifically through the maternal strain, with CAST, PWK and WSB having negative “dam” effects on litter size (Figure 3). It is likely that selection pressure in classical lab strains is associated with larger litters compared with the wild-derived strains. Additionally, two of the wild-derived strains, CAST and PWK, are from a different subspecific origin than the other six CC founders. This likely contributes to decreased productivity through subspecific incompatibilities (Shorter *et al.* 2017).

We identify and report environmental factors that may influence litter size. The breeding of this diallel was performed across two different vivariums over the course of 4 years, and we see a significant effect from housing facility. The Hillsborough facility was associated with larger litters for all strains, especially 129S1 and WSB. The two facilities have many different factors that could explain these differences. The Hillsborough facility housed multiple species, including dogs and mice, had smaller rooms that held approximately 300 mouse cages, was remotely located in a rural area, had different laboratory personnel, had cage changes once a week, and was supplied with filtered well water. The diallel breeding at the GMB facility took place in a large central room, contained only laboratory mice, is located in a basement of a large seven story research building on campus, has cage changes every other week, and is supplied with filtered city water. It is possible that one or more of these factors, independently or in combination, affected productivity. Another finding was that seasonality, which has previously been shown to influence litter size and frequency of litters in mammals (Drickamer 1990), did not seem to significantly impact litter size in this study. This is likely due to consistent light-dark cycles and temperatures as well as a steady diet. We did observe a significant effect on litter size after the transfer of the mice to the UNC GMB facility (February 2010), which reduced overall litter sizes from March to June 2010. This may have been due to the use of ivermectin during the time of the transfer. Other factors, such as the sex of the laboratory personnel interacting with the animals, are generally known to influence rodent behavior and could contribute to some of the environmentally-induced variation we observed(Sorge *et al.* 2014).

Last, we measured sex ratio across all inbred and outbred crosses. Despite some departures from equality at the nominal significance threshold (alpha = 0.05), no associations with founder strain dosage remain significant after correction for multiple testing. Recent work has suggested a potential for bias in sex ratio driven by the male germline in *Mus musculus* (Conway *et al.* 1994; Macholán *et al.* 2008; Cocquet *et al.* 2009; Ellis *et al.* 2011; Cocquet *et al.* 2012; Turner *et al.* 2012; Larson *et al.* 2016), particularly in intersubspecific hybrids that are mismatched for copy number of X- and Y-linked genes expressed in postmeiotic spermatids. Although there are no previous observations that suggest a bias in sex ratio in the CC or its founders, it remains an important characteristic to measure in a study on reproductive productivity.

To estimate heritable effects on liter size and sex ratio we extended the original BayesDiallel model of Lenarcic *et al.* (2012) in two new ways. First, to better understand and distinguish the effects arising from female and male parents, we reparameterized our strain-specific additive and parental-sex effects such that we could provide estimates of maternal strain and paternal strain effects separately. In the original BayesDiallel model, maternal and paternal strain effects are split into “additive” effects, which consolidates the effects they have in common, and “parental sex”, which models any remaining deviation between the two. Recognizing that additive effects from the sire are essentially wiped out by the additive effects, we instead collapsed the additive and parental-sex effects into “dam.strain” and “sire.strain” effects in a post-processing step on the posterior output. This allowed us to run the original BayesDiallel model while also viewing our data from the perspective of dam strain and sire strain contributions.

Second, we reimplemented the original MCMC sampler, designed for modeling a continuous outcome variable, in a general package MCMCglmm (Hadfield 2010) in order to model count and binary responses. Litter size, as measured, is most naturally distributed as Poisson, with zero-truncation owing to the fact that only successful litters were recorded. Sex ratio is most naturally modeled as a binomial, with an underlying (male) proportion between 0 and 1. Although it would be possible to obtain an approximate analysis by transformations to normality using the original BayesDiallel (and we do this for litter size for some otherwise hard-to-obtain quantities), we found such approximations to be inadequate for reliable estimation of higher order effects in the case of litter size and deeply flawed in the case of sex ratio.

Although there is a computational cost, and added complexity to determining variance contributions, this new implementation achieves several objectives: 1) we no longer break the assumptions in the original model regarding normally distributed errors; 2) we easily accommodate overdispersion in our data; and 3) we can select from a large number of GLMMs ones that more closely resemble the forms of our data observations. In addition, we believe this flexibility will be appealing to many other researchers who would like to model non-Gaussian distributed phenotypes using diallel designs, and we have provided the code in an R package litterDiallel (https://doi.org/10.5281/zenodo.1435829).

Overall, these results have implications for other avenues of future research. Future multiparental research populations should test for strain incompatibilities, reproductive phenotyping, and other health traits in a full diallel before the recombinant inbreeding begins (Odet *et al.* 2015). These future research populations should also use non-related wild-derived individuals from the same subspecific origin in order to increase genetic diversity without introducing hybrid incompatibilities.

## Acknowledgments

This work was supported in part by the following grants from the National Institutes of Health: P50GM076468, P50HG006582/P50MH090338, R01HD065024 (FP-MV), T32HD040127 (JRS), T32AI007419 (PLM), R01GM104125 (PLM), and R35GM127000 (WV). The Collaborative Cross project is also supported by the University Cancer Research Funds granted to Lineberger Comprehensive Cancer Center (MCR012CCRI).

## Appendix A: Diallel Model For Litter Size

We collected data on litter size at weaning for 62 genetic crosses of inbred lines, across four years of breeding.

We use zero-truncated poisson (ZTP) regression for modeling our data. This type of regression is explicit in its framework accounting for discrete observations, flexible in its ability to use linear mixed models on the latent scale, and allows for parameterization of excess variance observed, in a way that standard Poisson regression does not. We account for the zero depletion in our data by using ZTP regression instead of standard Poisson regression, since we exclude observations of birth cohorts where no pups survived to weaning.

In Figure 4, the distribution of the observed data (litter size) is displayed for the WSBxWSB inbred mating, along with simulated data from a zero-truncated Poisson distribution based on the data mean.

**Figure 4.**
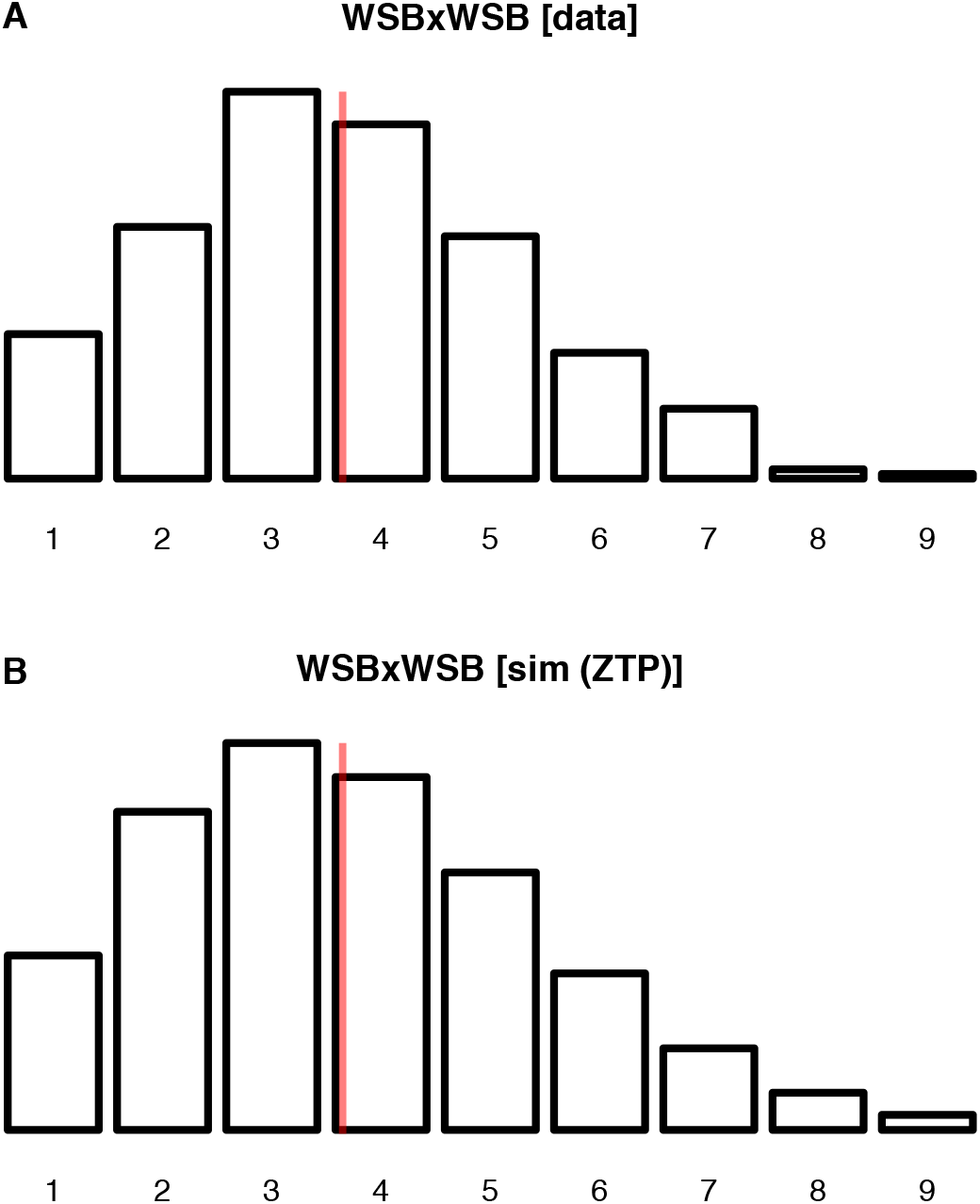
Comparison of observed data and (pseudo-)randomly generated data for the WSBxWSB inbreds. (A) Distribution of the litter diallel data for WSBxWSB (nlitters=341, npups=1244). (B) Frequencies expected from a zero-truncated Poisson distribution with mean=3.648 (same as in A). The vertical red lines indicate the mean value.

For the ZTP, the first two moments (mean and variance), for values *y_i_* > 0, are given by:

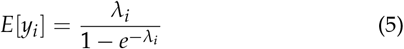

The relationship between the standard Poisson regression model (with overdispersion) is shown in Figure 5. The ZTP modifies this by truncating values on the expected (log) data scale and observed data scale to be 1 or greater.

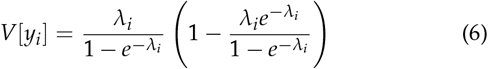

**Figure 5.**
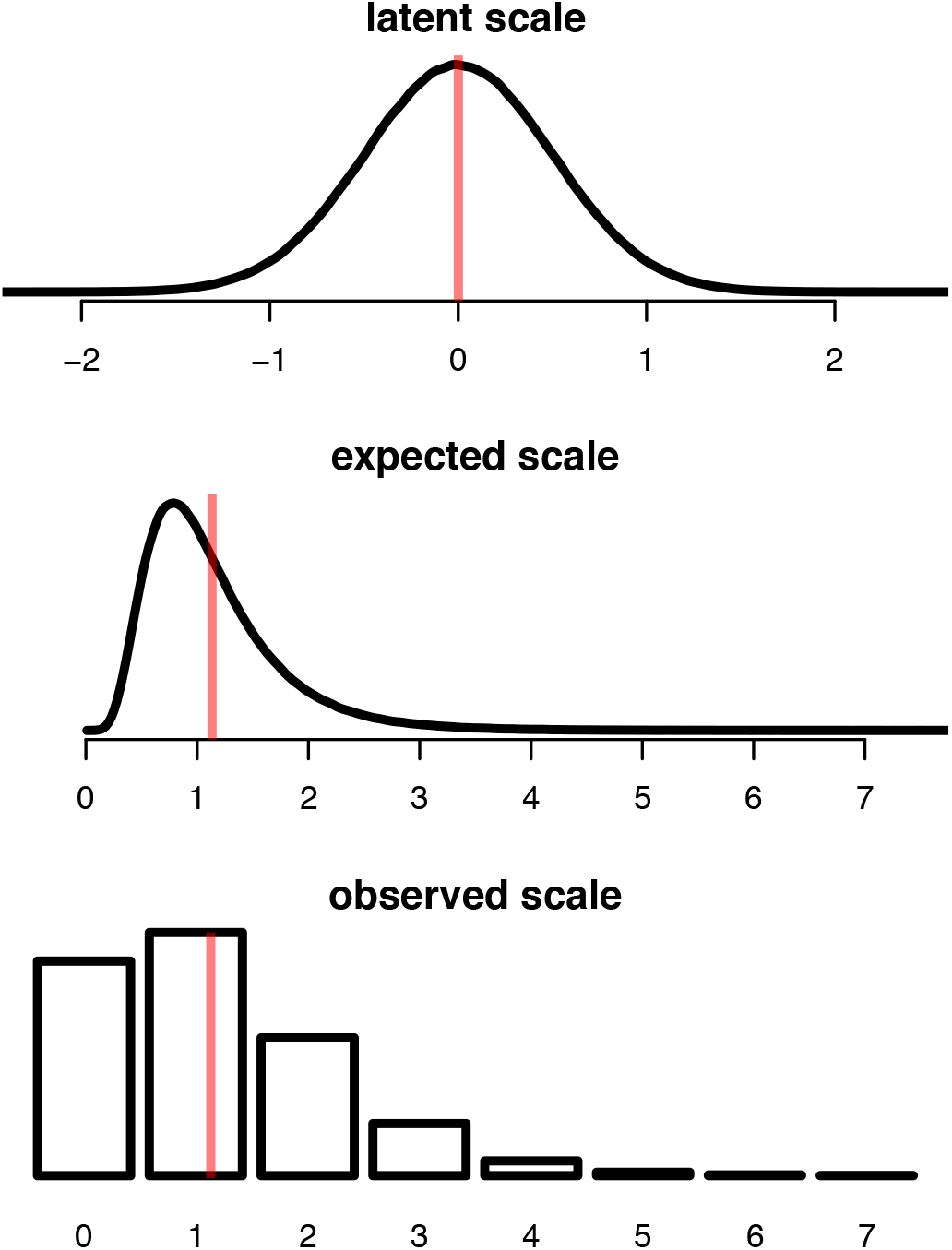
Latent and data scale comparison. A toy example illustrating standard Poisson regression of count data, with overdispersion, giving values drawn from pseudorandom normal variables drawn from *?(μ* = 0, sd = 0.5), showing how the continuous *latent* scale values [top], which correspond to the scale of the linear predictor, map onto their *expected* (data) scale value (through the inverse link, *f* (x) = *e*^x^) [middle], and how the mean *expected* (data) scale value corresponds to the *observed* (data) scale values (integers). In each frame, the vertical red line corresponds to the mean of the values shown in that frame.

### Diallel effect estimates

Diallel effect estimates are obtained using an MCMCglmm (Hadfield 2010) implementation of BayesDiallel (BayesDiallel-glmm, henceforth), with 1.515 x 10^6^ iterations, 1.5 x 10^4^ iterations of burn-in, and thinning by 500 (saving only every 1/500^th^ iteration), to obtain 3000 independent samples with minimal autocorrelation, and plotted as highest posterior densities (HPDs) in the results.

### Conversion to the expected data scale

The effects estimated from the BayesDiallel-glmm model are transformed from the latent scale to the (expected) data scale via the inverse link function, *i.e.* exp(a_AJ_), for interpretability of effects on the original data scale.

### Variance Explained Using Variance Projection

To avoid the problem of interpretability in transforming variance parameter (and variance projection) estimates from the latent to the observed data scales, we instead calculate and report variance projections, as calculated on the Gaussian version of BayesDiallel. In order to account for heteroscedasticity (unequal variance) of the model residuals that arises from the approximately ZTPoisson nature of the data, we use a variance-stabilizing transformation (VST) (Yu 2009) and run BayesDiallel again (Gaussian) to obtain Variance Projections on the modeled data. The VarPs that are calculated from these parameter estimates are an approximation of the variance contributions that we would observe in the GLMM BayesDiallel model.

## Appendix B: Diallel Model For Sex Ratio

We model the male pup counts and female pup counts jointly, using the BayesDiallel linear model, formulated for binomial GLMM regression. This model directly considers genetic effects on the imbalance in male vs. female pups by parameterizing the number of males and number of total pups (or the total, and the fraction of males in the total). The model is elaborated in the methods section of the main manuscript.

The proportion of weaned pups that are male, or equivalently, the proportion of weaned pups that are female, is approximated by a binomial distribution. In our data, the mean and the variance of male ratio are 0.494 and 0.063, respectively.

To generate the upper and lower 95% boundaries, as shown in Figure S6, for the expected phenotype under the null hypothesis of male pup proportion = 0.5, we used the qbinom function in the stats package in R.

